# Plasma and Fecal Metabolite Profiles in Autism Spectrum Disorder

**DOI:** 10.1101/2020.05.17.098806

**Authors:** Brittany D. Needham, Mark D. Adame, Gloria Serena, Destanie R. Rose, Gregory M. Preston, Mary C. Conrad, A. Stewart Campbell, David H. Donabedian, Alessio Fasano, Paul Ashwood, Sarkis K. Mazmanian

## Abstract

Autism Spectrum Disorder (ASD) is a neurodevelopmental condition with hallmark behavioral manifestations including impaired social communication and restricted repetitive behavior. In addition, many affected individuals display metabolic imbalances, immune dysregulation, gastrointestinal (GI) dysfunction, and altered gut microbiome compositions. We sought to better understand non-behavioral features of ASD by determining molecular signatures in peripheral tissues. Herein, we present the untargeted metabolome of 231 plasma and 97 fecal samples from a large cohort of children with ASD and typically developing (TD) controls. Differences in lipid, amino acid, and xenobiotic metabolism discriminate ASD and TD samples. We reveal correlations between specific metabolite profiles and clinical behavior scores, and identify metabolites particularly associated with GI dysfunction in ASD. These findings support a connection between GI physiology, metabolism, and complex behavioral traits, and may advance discovery and development of molecular biomarkers for ASD.

## Introduction

Many diseases are associated with informative metabolic signatures, or biomarkers, that enable diagnoses, predict disease course, and guide treatment strategies. In contrast, autism spectrum disorder (ASD) is diagnosed based on observational evaluation of behavioral symptoms, including reduced social interaction and repetitive/stereotyped behaviors (*1*). The average age of ASD diagnosis is between 3-4 years old (*2*), at which time children can receive behavioral therapy, the gold standard treatment. Because earlier diagnosis improves efficacy of behavioral therapies (*3, 4*), molecular biomarkers represent an attractive approach for identifying ‘at-risk’ populations and may aid development of personalized therapies. This prospect is increasingly important given the rising rate of ASD diagnoses, which currently stands at up to 1 in 59 children in the United States (*2*), with no FDA approved drugs for core behavioral symptoms.

Metabolic abnormalities have been reported in ASD (*5*), though most have measured only a small subset of metabolites and many outcomes have not reproduced between cohorts. Mitochondrial disease, which heavily influences systemic metabolism, is estimated to be higher in ASD compared to controls (5% vs ~0.01%) (*6*), and genes crucial for mitochondrial function are risk factors for ASD in humans and rodent models (*7*). The metabolic abnormalities associated with mitochondrial dysfunction in ASD affect cellular energy, oxidative stress, and neurotransmission in the gut and the brain (*7–19*). Other metabolic profiles in ASD implicate aromatic and phenolic metabolites, including derivatives of nicotinic, amino acid, and hippurate metabolism (*20–30*). Various amino acids are detected at differential levels across studies and across sample types, but any consistent patterns are difficult to discern (*15, 24, 26, 29, 31–34*).

Some of the discrepancies between these studies are likely due to differences in sample number, tissue analyzed, and methodology. Other sources of variability include differences in environmental factors, such as diet and gut bacteria, which differ between ASD and TD populations (*35–37*). Diet is a major source of circulating metabolites, impacting the metabolome directly or indirectly through chemical transformation by the trillions of gut microbes, the microbiome, which has been proposed to modulate complex behaviors in animal models and humans (*38–40*). Such proposed environmental modulators of ASD may integrate with genetic risks to impact behavioral endpoints through the actions of small molecules produced in peripheral tissues outside the brain.

Herein, we present a comprehensive comparison of an extensive panel of identified metabolites in human plasma and feces from a large cohort of matched ASD and TD children. We identified differential levels of metabolites ranging from hormones, amino acids, xenobiotics, and lipids, many of which correlate with clinical behavior and GI scores. To our knowledge, this is the first study to concurrently evaluate paired intestinal and systemic metabolomes in a high-powered analysis with a large number of identified metabolites, allowing direct associations between metabolites previously highlighted in ASD samples and discovering new metabolites of interest. These findings support the emerging concept of evaluating non-behavioral features in the diagnosis of ASD and its GI comorbidities.

## Methods And Materials

### Participants

Samples for this study, aged 3-12 years old, were collected through the UC Davis MIND institute (*41, 42*). ASD diagnosis was confirmed at the MIND Institute by trained staff using the Autism Diagnostic Observation Schedule (ADOS), and the Autism Diagnostic Interview-Revised (ADI-R). Subjects were diagnosed prior to 2013 based on DSM IV. Typically developing (TD) participants were screened using the Social Communication Questionnaire (SCQ). Participants in the TD group scored within the typical range, i.e. below 15, on the SCQ and above 70 on the Mullen Scales of Early Learning (MSEL) and Vineland Adaptive Behavior Score (VABS). Ninety-seven of the participants, who also provided the stool samples, completed an additional evaluation to determine gastrointestinal (GI) symptoms. GI status was determined using the GI symptom survey (GISS), based upon Rome III Diagnostic Questionnaire for the Pediatric Functional GI Disorders (*43*) and described in detail elsewhere (*42*).

See Supplemental materials for additional methods.

## Results

### Plasma and Fecal Metabolomes Differ between ASD and TD

Plasma samples from 130 ASD and 101 TD children were analyzed along with fecal samples collected from a subset of these same ASD (n=57) and TD individuals (n=40) (Figures S1A–S1G). Samples and metrics of behavioral and GI scores were obtained from the UC Davis MIND institute (*41, 42*). The ASD group was stratified into subsets of children with GI symptoms (ASD+GI) or without GI symptoms (ASD-GI), to explore potential effects of comorbid intestinal dysfunction in ASD (40 out of 130 ASD samples were +GI). This stratification was based on symptoms associated with ASD including diarrhea, constipation, and irritable bowel syndrome-like symptoms.

Samples were analyzed by the global metabolite panel (plasma and fecal samples) and the complex lipid panel (plasma samples) by Metabolon, Inc. (Durham, NC), which identified a total panel of 1,611 plasma and 814 fecal metabolites (Tables S1-S2). Overall, we discovered that 310 metabolites were differentially abundant between the ASD and TD groups in plasma and 112 metabolites were differentially abundant in feces (Figures 1A-1B, S1C, and Tables S1-S2). Using a quantitative assay for a targeted panel of metabolites, we observed a high correlation between relative abundance and precise concentration (Figure S2A). Overall, these data expand on previous evidence that the metabolomic profile of ASD and TD populations display differences not only in the gut compartment, but also in circulation, which may affect the levels of metabolites throughout the body, including the brain (*44, 45*).

**Figure 1.**
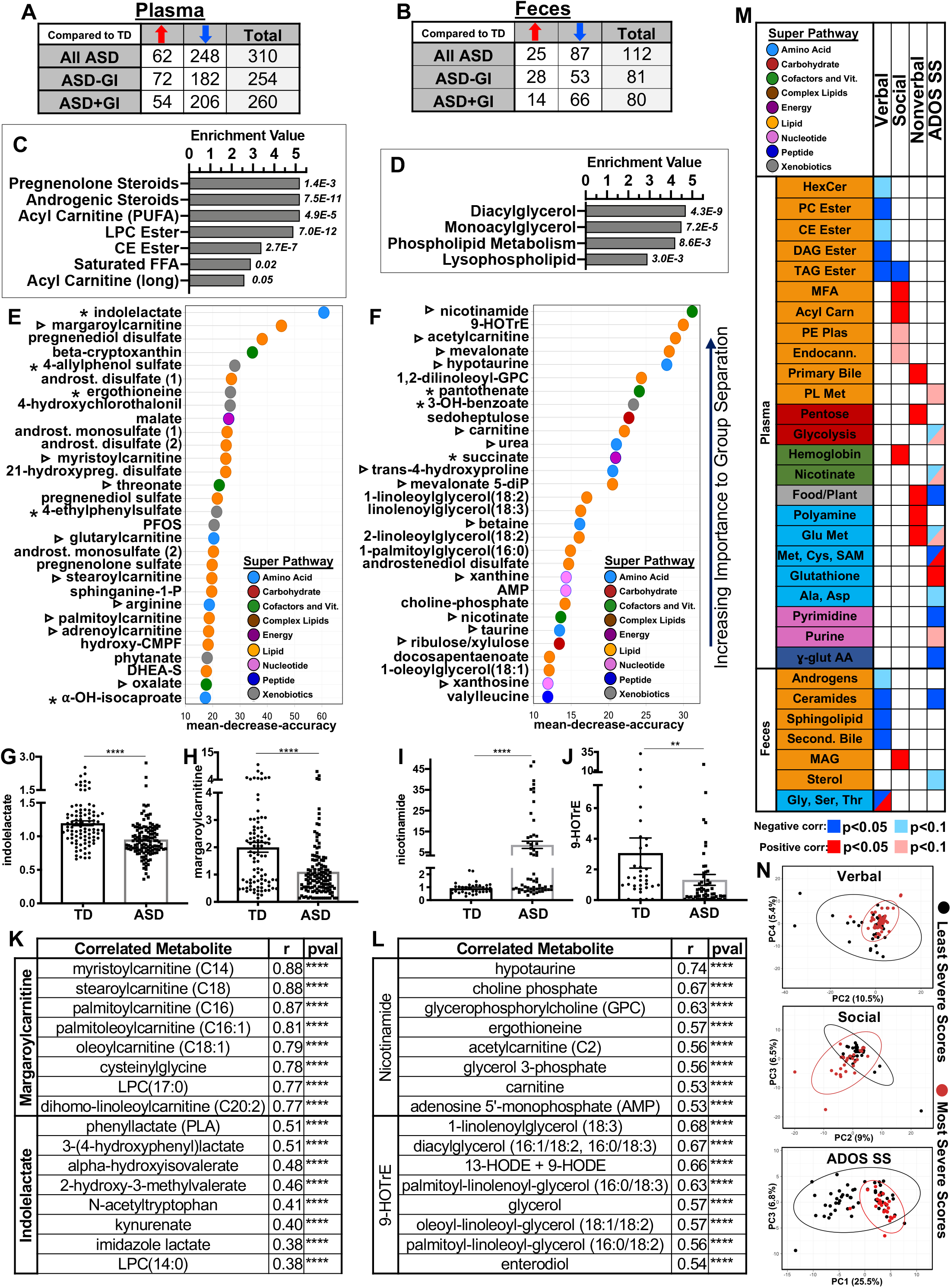
Plasma and Fecal Metabolomes Differ between ASD and TD and Global Metabolite Levels Correlate with Clinical Behavioral Scores. (A-B) The number of significantly elevated and decreased metabolites (pval<0.05) in ASD samples compared to the TD control group by ANOVA contrasts in plasma and feces, respectively. Samples are stratified by all samples or samples without or with GI symptoms (-GI, +GI). *See also Table S1 and S2*. (C-D) Pathway analysis results of human plasma and fecal comparisons (all samples), indicating which metabolomic pathways are the most altered between groups, with enrichment value plotted and p-value to the right of each bar. Metabolites within each pathway could be observed at either higher or lower levels, as this plot only indicates significant changes. (E-F) Top 30 most distinguishing metabolites between each group in plasma and feces by random forest analysis, with mean decrease accuracy along the x-axis. Metabolites known to be produced by (asterisks) or influenced by (triangles) the gut microbiota are denoted. The super pathway to which each metabolite belongs to is indicated by color of sphere and defined in the legend. *See also Figure S1*. (G-H) Scaled intensity values indicating relative levels of the most distinguishing molecules between ASD and TD (all samples) in plasma. Asterisks indicate significance in ANOVA contrasts performed on total metabolomics dataset. p values: 2.87E-09(G) and 1.57E-06(H). Data are represented as mean ± SEM. (I-J) Scaled intensity values indicating relative levels of the most distinguishing molecules between ASD and TD (all samples) in feces. Asterisks indicate significance in ANOVA contrasts performed on total metabolomics dataset. p values: 1.37E-05(I) and 0.002(J). Data are represented as mean ± SEM. (K) Top correlated plasma metabolites that covary with margaroylcarnitine and indolelactate. All p-values <0.0001. (L) Top correlated fecal metabolites that covary with nicotinamide and 9-HOTrE. All p-values <0.0001. (M) Spearman correlations between behavior scores of ASD children in the ADIR diagnostic test (Verbal, Social, and Nonverbal metrics) and the ADOS-SS and metabolite pathways in ASD samples. Directionality of correlation is indicated in the legend at bottom. Colors of pathways are defined at the top left of the chart. A split box means that both positive and negative correlations occur with metabolites within that pathway. *See also Tables S1 and S2*. (N) PCA plot comparing the metabolic profile least and most severe ~quartiles within verbal, social and ADOS-SS scores. PCA input included all metabolites significantly associated with the behavior. Clustering is denoted by ellipses of the 95% confidence interval. LPC, lysophosphatidylcholine; CE, cholesterol ester; FFA, free fatty acid; androst., androstane; hydroxypreg, hydroxypregnenalone; PFOS, perfluorooctanesulfonic acid; hydroxy-CMPF, hydroxy-3-carboxy-4-methyl-5-propyl-2-furanpropionate; DHEA-s, dehydroepiandrosterone sulfate; 9-HOTrE, 9S-hydroxy-10E,12Z,15Z-octadecatrienoic acid; AMP, adenosine monophosphate; HExCer, Hexosylceramide; PC, phosphatidylcholine; DAG, diacylglycerol; TAG, triacylglycerol; MFA, monounsaturated fatty acid; PE plas, phosphatidylethanolamine plasmalogens; Endocann, endocannabinoid; Met, methionine; Cys, cysteine; SAM, s-adenosyl methionine; Ala, alanine; Asp, aspartate; γ-glutAA, gamma-glutamyl amino acids; Second. Bile, secondary bile; MAG, monoacylglycerol; Gly, glycine; Ser, serine; thr, threonine. *See also Figures S1 and S2*.

To appreciate the biological relevance of different metabolomes between ASD and TD, individual metabolites were integrated into biochemical pathways for pathway enrichment analysis, revealing the degree of change within each. Here, we identified large scale changes, mostly in lipid, xenobiotic, and nucleotide pathways associated with diverse physiological processes (Figures 1C-1D, S2B). We then used Random Forest machine learning analysis to determine if metabolite profiles can unbiasedly predict whether the sample came from an ASD and TD donor. Overall, the modest predictive accuracy of this machine learning approach was 68% for plasma and 67% for feces. To test whether focusing on the most discriminating metabolites would improve the prediction, we repeated the Random Forest analysis using the top 30 metabolites and found that the predictive accuracy for plasma improved slightly to 69% when using all ASD samples, and to 73% when using only ASD-GI samples. For fecal samples, the predictive accuracy improved to 75% using all ASD and to 73% using only ASD-GI. The top 30 metabolites, calculated by measuring the mean decrease in accuracy of the machine learning algorithm, are useful to describe the strongest drivers of overall metabolic differences between ASD and TD populations. These most discriminatory metabolites were primarily from the lipid, amino acid, xenobiotic, and cofactor/vitamin super pathways (Figures 1E–1F). Several of these metabolites have been previously linked to ASD, such as steroids, bile acids, acyl-carnitines, and nicotinamide metabolites (*46–49*). Further, multiple molecules known to be produced or manipulated by the gut microbiota also featured prominently, including 4-ethylphenyl sulfate (4EPS), which is elevated in an ASD mouse model, and indolelactate, a microbe-derived tryptophan metabolite (Figure 1E) (*50, 51*). The two most discriminatory molecules in plasma (Figures 1G–1H) and feces (Figures 1I–1J) are depicted. Metabolites correlating the strongest with these discriminatory metabolites are closely related on a structural and metabolic level (Figures 1K–1L).

### Global Metabolite Levels Correlate with Clinical Behavioral Scores

Using clinical metadata for ASD individuals, we correlated the levels of individual metabolites to the verbal, social, and nonverbal scores of standard diagnostic measures: the Autism Diagnostic Interview, Revised (ADIR), a parent questionnaire, and the cumulative Autism Diagnostic Observation Schedule severity score (ADOS-SS), conducted by trained health professionals (*1*) (Tables S1-S2). Next, grouped correlations between the behavior metrics and entire metabolite pathways were calculated (Figure 1M). We found that verbal and social scores primarily correlate with lipid metabolism pathways and that nonverbal scores have the fewest correlations. The ADOS-SS correlated with diverse metabolite pathways, including amino acids and food/plant component pathways, which may be partly due to the diverse array of symptoms integrated into the ADOS-SS score. Additionally, when stratified by severity of behavioral scores, we observe modest clustering according to plasma metabolites correlated to the disorder between the top and bottom quartiles of behavior severity groups for verbal, social, and ADOSS-SS metrics, but not for nonverbal, and less so with fecal samples (Figures 1N, S1H–S1I). These correlations support the involvement of lipid, amino acid, and xenobiotic metabolism in the etiology of ASD, as previously described (*8, 48, 52*), and reveal new candidates for ASD biomarkers that correlate with symptom severity.

### Transfer of ASD Fecal Microbiota into Mice is Accompanied by Metabolic Signatures

Since microbial metabolites ranked highly in the Random Forest machine learning analysis, we tested if any of the observed metabolite differences in humans could be transferred to mice via fecal microbial transplant. We selected 4 male donor samples from each of the ASD and TD groups and colonized 2-3 male germ-free mice per donor for three weeks before collecting plasma and fecal samples for metabolite profiling and bacterial DNA sequencing (Figure S2C–S2D), respectively. Global metabolomic analysis revealed that colonized mice modestly cluster by donor and group when statistically significant metabolites are considered in PCA analysis (Figure S2E). We selected the human donors based on 4EPS levels (Figure S2F), due to its involvement in an ASD mouse model (*50*) and dysregulation of similar phenolic compounds in human ASD (*22, 24, 28, 29, 53*). 4EPS is not produced by the host and is strictly a bacterial metabolite (*50, 54*). Surprisingly, we observed 4EPS levels in a bimodal distribution in mouse samples (Figure S2G–S2H). In spite of the surprising results with 4EPS, many of the metabolites with the highest fold change and lowest *p*-value are indeed other phenolic molecules such as metabolites of hippurate, tyrosine, and diet-derived phenols (Figures S2I, Table S5). While preliminary, these studies reveal microbiome-mediated effects on xenobiotic pathways and phenolic metabolites dysregulated in ASD.

### Lipid and Xenobiotic Metabolites Correlate with GI Symptoms

Considering the prevalence of intestinal issues in ASD, GI-dependent metabolite analysis between individuals may provide useful insights. Accordingly, we found a total of 87 and 24 metabolites in plasma and feces, respectively, that were differentially abundant within the ASD+GI compared to ASD-GI individuals (Figures 2A–2B). At the pathway level, we found differences in free fatty acid and xenobiotic metabolites in the food component and plant pathways between ASD+GI vs ASD-GI plasma (Figure 2C), with no broad GI-dependent pathway alterations in fecal samples (Table S2). In plasma, free fatty acids of multiple chain lengths were lower in ASD+GI compared to ASD-GI samples, including monounsaturated, saturated, and polyunsaturated fatty acids (PUFAs) (Figure 2D). PUFAs are anti-inflammatory and lower levels of these fatty acids may contribute to GI dysfunction directly (*12*).

**Figure 2.**
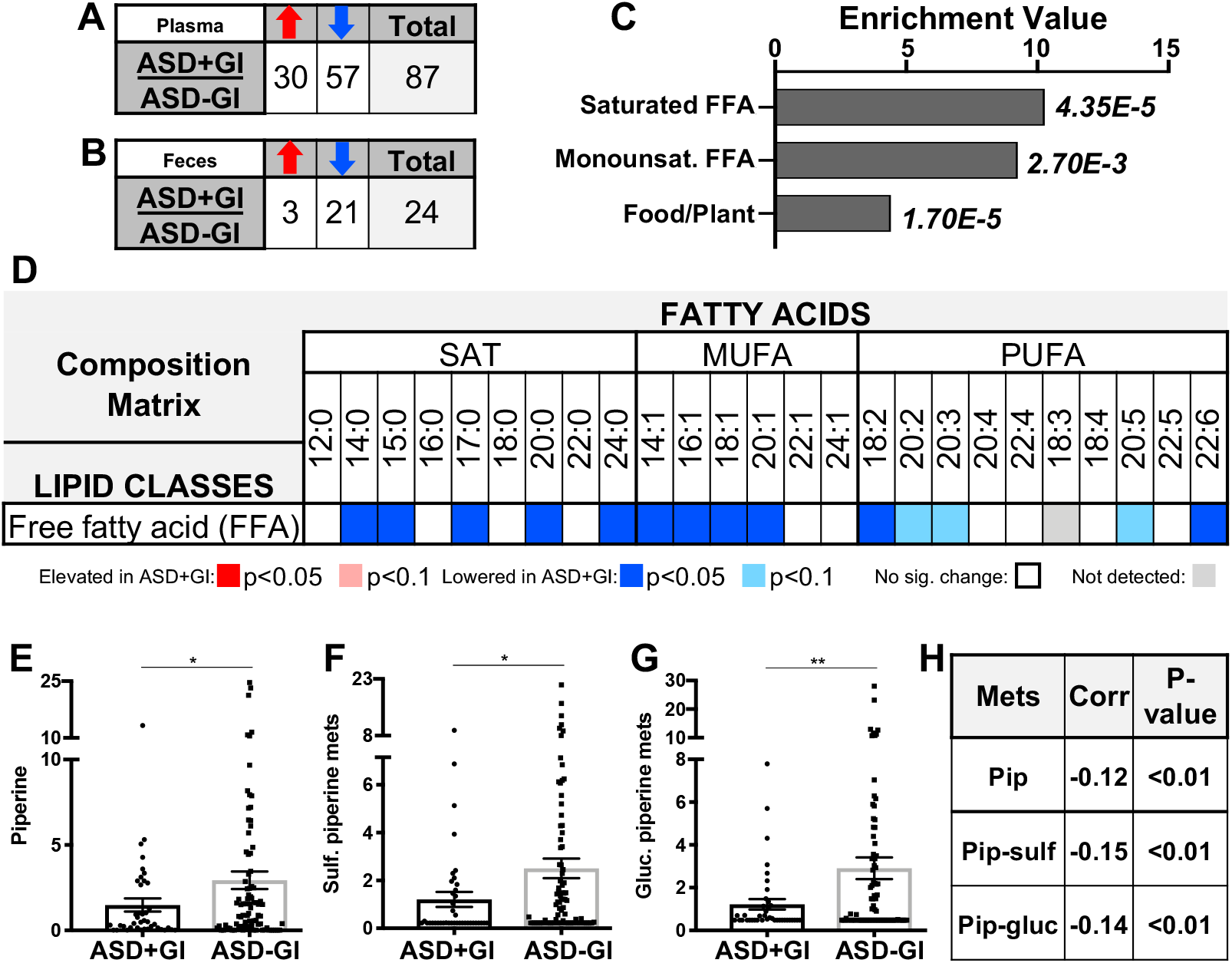
Lipid and Xenobiotic Metabolites Correlate with GI Symptoms. (A-B) Numbers of significantly (p value<0.05) elevated and decreased metabolites within the ASD samples, comparing ASD+GI to ASD-GI in plasma and feces, respectively, by ANOVA contrasts. (C) All pathways significantly altered in the comparison between ASD+GI and ASD-GI human plasma samples, with enrichment value plotted and p-value to the right of each bar. Metabolites within each pathway could be observed at either higher or lower levels, as this plot only indicates significant changes. (D) Complex lipid panel results for the ASD+GI vs ASD-GI plasma samples. Along the top, length of lipid chain of the free fatty acid is displayed (chain length:number of unsaturations) and are grouped into saturated (SAT), monounsaturated (MUFA), and polyunsaturated (PUFA) fatty acids. Color legend is displayed below. (E-G) Scaled intensity values indicating relative levels of free and conjugated piperine in ASD+GI and ASD-GI symptoms. Data are represented as mean ± SEM. Asterisks indicate significance in unpaired t test with Welch’s correction. P-values: 0.03(E), 0.01(F), and 0.003(G). Free piperine does not reach statistical significance in ANOVA comparison performed on total dataset (p value=0.16). Individual p values (all<0.05) of each sulfated and glucuronidated form in ANOVA comparison can be found in *Table S1*. (H) Spearman correlation of piperine and derivatives with ADOS-SS. *See also Figure S1*. FFA, free fatty acid; monounsat., monounsaturated.

Several metabolites from the food component and plant pathway also discriminate ASD+GI from ASD-GI, such as lower circulating piperine metabolites in ASD+GI plasma samples (Figures 2E-2G). A lower level of piperine metabolites was also associated with higher ADOS-SS (Figure 2H) and worse nonverbal scores (Table S1). Observed alterations to piperine metabolite levels are supported by the fact that oral administration of piperine has been successfully used as a treatment in preclinical ASD models, presumably due to its antioxidant quality (*55*). While these correlations are interesting, the implications of correlating an altered metabolome and GI symptoms in ASD remain to be determined.

### Steroid Hormone Levels are Elevated in ASD

Multiple human ASD studies have examined the levels of specific steroid metabolites within androgenic, pregnenolone and progesterone metabolism, with some finding aberrant levels, positive correlations with ASD severity, and behavioral improvement following treatments that lower levels of certain hormones (*47, 56–64*). On the other hand, in a recent clinical trial of ASD children given an antioxidant treatment, levels of pregnenolones and androgens increased and correlated with improved behavior (*65*). In our dataset, we found robust increases of almost every detected metabolite within the pregnenolone, androgen, progesterone, and corticosteroid pathways in the plasma of ASD children (Figures 1C, 3A, Table S1). Similarly, some androgenic steroid pathway metabolites were elevated in ASD fecal samples (Figures 3A, Table S2). This is a strong indicator that the physiological pathways associated with the downstream metabolism of cholesterol are significantly altered between ASD and TD populations (Figure 3B). There does not appear to be a global change in steroid metabolism, as most primary bile acid and sterol metabolites were unaffected (Tables S1-S2). We observed some elevation of these hormone levels independent of sex, which is notable considering the male bias in ASD, reflected in our primarily male sample set (7-15% female) (*66*) (Figure S3A–S3B). Because a cluster of our samples are from older individuals in the ASD group, and to account for age-dependent increases in androgens, we stratified by age and still observed heightened androgenic and pregnenolone metabolite levels in ASD subpopulations (Figure S3C). Taken together, these data indicate that steroidal hormone metabolism may be altered in the ASD population relative to TD samples and that these differences are not driven solely by sex or age differences in our cohort.

**Figure 3.**
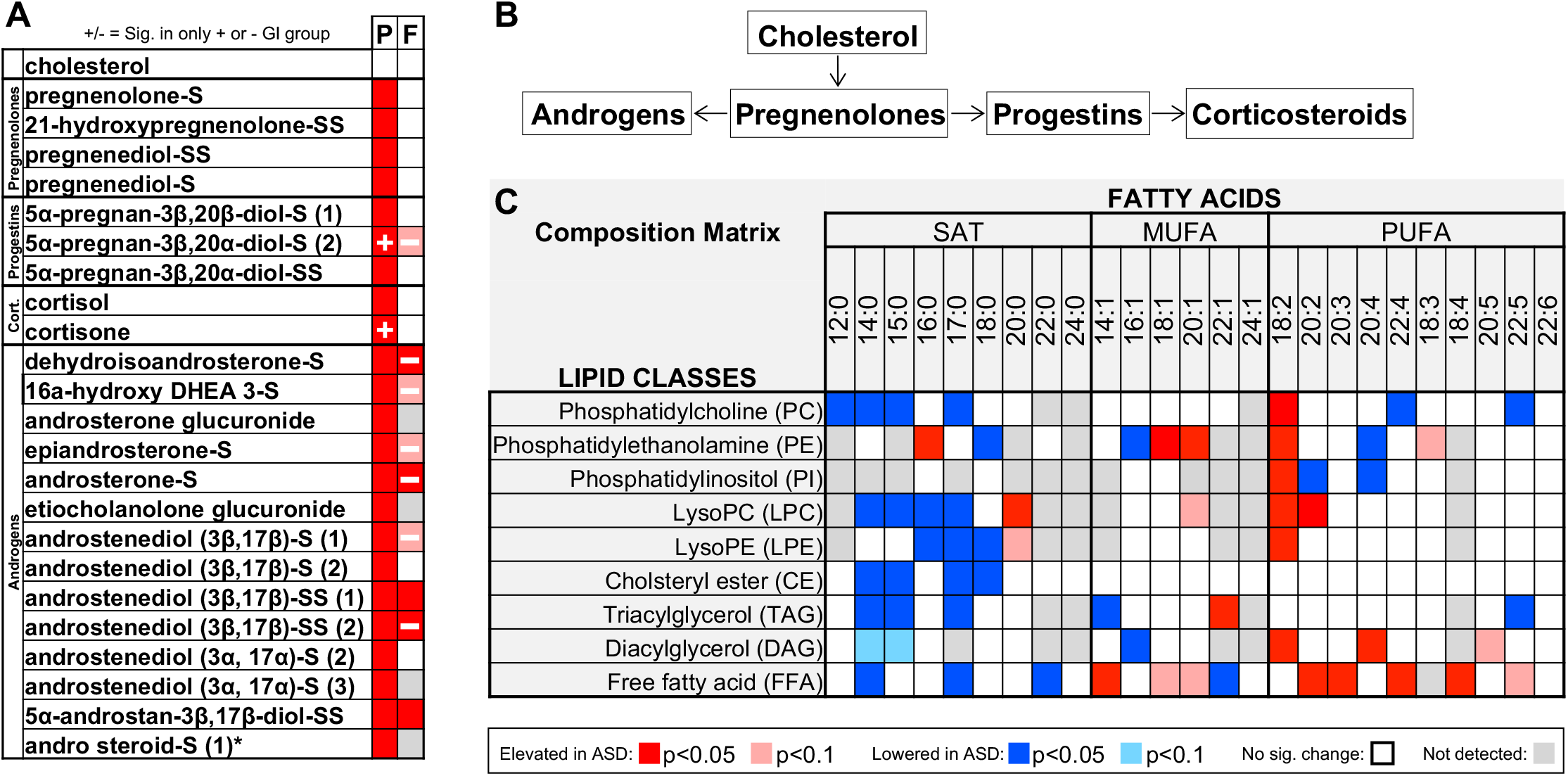
Steroid Hormone Levels are Elevated in ASD and other Lipid Metabolite Levels Differ in ASD. (A) Significant alterations to levels of all metabolites detected in the pregnenolone, progestin, and androgen steroid pathways in plasma (P) and feces (F), with colors indicating significance and fold change according to legend. (B) Schematic summarizing the metabolism of these pathways. (C) Complex lipid panel results for all the ASD plasma samples compared to TD controls with acyl chain length of lipids across the top, described by chain length, degree of unsaturation and categorized by saturated (SAT), monounsaturated (MUFA), and polyunsaturated (PUFA) fatty acids. Lipid classes are listed along the left. Direction of change and significance are indicated by the legend. Significance determined by ANOVA contrasts. *See also Figure S3 and Table S3*.

### Lipid Metabolite Levels Differ in ASD

Lipids are crucial for energy storage, cellular membrane integrity and cell signaling. They play a variety of roles in the central nervous system, and their dysregulation has been linked to ASD (*9, 11*). Phospholipids have been measured at lower levels, while long chain fatty acids are reportedly elevated but PUFAs have been measured at both higher and lower levels, depending on the cohort (*9, 12, 13*). Here, we performed an untargeted, quantitative metabolite analysis on complex lipids. The concentrations of 999 lipids were quantified, and 13.7% of these were significantly different in the ASD samples, with 4.6% increased and 9.1% decreased compared to TD samples. Many of these differentially abundant lipids included phospholipids, cholesterol esters and glycerolipids. In general, shorter (14-18 chain length) saturated fatty acids were less abundant in ASD throughout the lipid classes (Figures 3C, S4A–S4B).

PUFA lipid levels were also different, with elevations in diacylglycerols and free fatty acids, and an enrichment of the 18:2 (linolenic) chain length in most lipid classes (Figure 3C). Fecal samples generally trended in the opposite direction from plasma in PUFA lipids (Figure S4C, Table S2). Intriguingly, multiple lipids with linolenic and linoleic (18:3) chains were correlated with social behavior (Tables S1 and S2). PUFA lipids containing linolenic and linoleic acids are precursors to the important PUFAs (arachidonic 20:4 and docosahexaenoic 22:6 acids) for brain development, function, and structural integrity (*12*). Future studies are needed to determine if these specific changes in lipid levels contribute to ASD symptoms.

### ASD Correlates with Cellular Energy and Oxidative Stress Metabolites

Many lipids are markers of mitochondrial and oxidative stress, offering a snapshot into cellular metabolic states. These markers include acyl-carnitines, which have been highlighted in various ASD studies and are established indicators of mitochondrial dysfunction (*11, 12, 14–18, 24, 25, 33, 46, 58, 67–71*). Acyl-carnitines are formed to allow transport of lipids across the mitochondrial membranes for beta-oxidation, and abnormal levels of these conjugated lipids accumulate to higher levels with a decrease in beta-oxidation. Interestingly, high levels of short acyl-carnitines are found in rodent models where ASD-like behaviors are induced with the short chain fatty acids, valproic and propionic acids (*72*).

We found differential levels of various acyl-carnitines in ASD, creating a pattern of more abundant short chain acyl-carnitines and less abundant long chain acyl-carnitines in the ASD-GI samples compared to TD samples (Figure 4A). Acyl-carnitines are positively correlated with more severe social defects, an effect driven by structures with shorter moieties (C2-C14) (Figure 1M, Table S1). In fecal samples, acetylcarnitine (C2) and free carnitine were elevated in ASD (Figures 4B–4C) and were highly discriminatory (Figure 1F). Other mitochondrial markers in both plasma and feces were also differentially abundant in ASD and are summarized in Figure 4D along with markers of phospholipid metabolism, which occurs largely in the mitochondria and was significantly altered in fecal samples (Figures 4D, 1D). Additionally, the 5 plasma metabolites most positively correlated with ADOS-SS are all involved in these cellular energy pathways (Figure S4D, Table S1). These observed differences in energy markers and lipids could have a neurodevelopmental effect during periods when the high lipid and energy requirement in the brain is crucial (*73–75*), and alterations to levels of tricarboxylic acid (TCA) cycle intermediates have been observed in human ASD prefrontal cortex samples (*49*). Such defects in cellular metabolism support the theory that mitochondrial dysfunction may not only be comorbid with ASD but also a potential contributing factor, as suggested by numerous previous reports (*6, 13, 67, 69, 72*).

**Figure 4.**
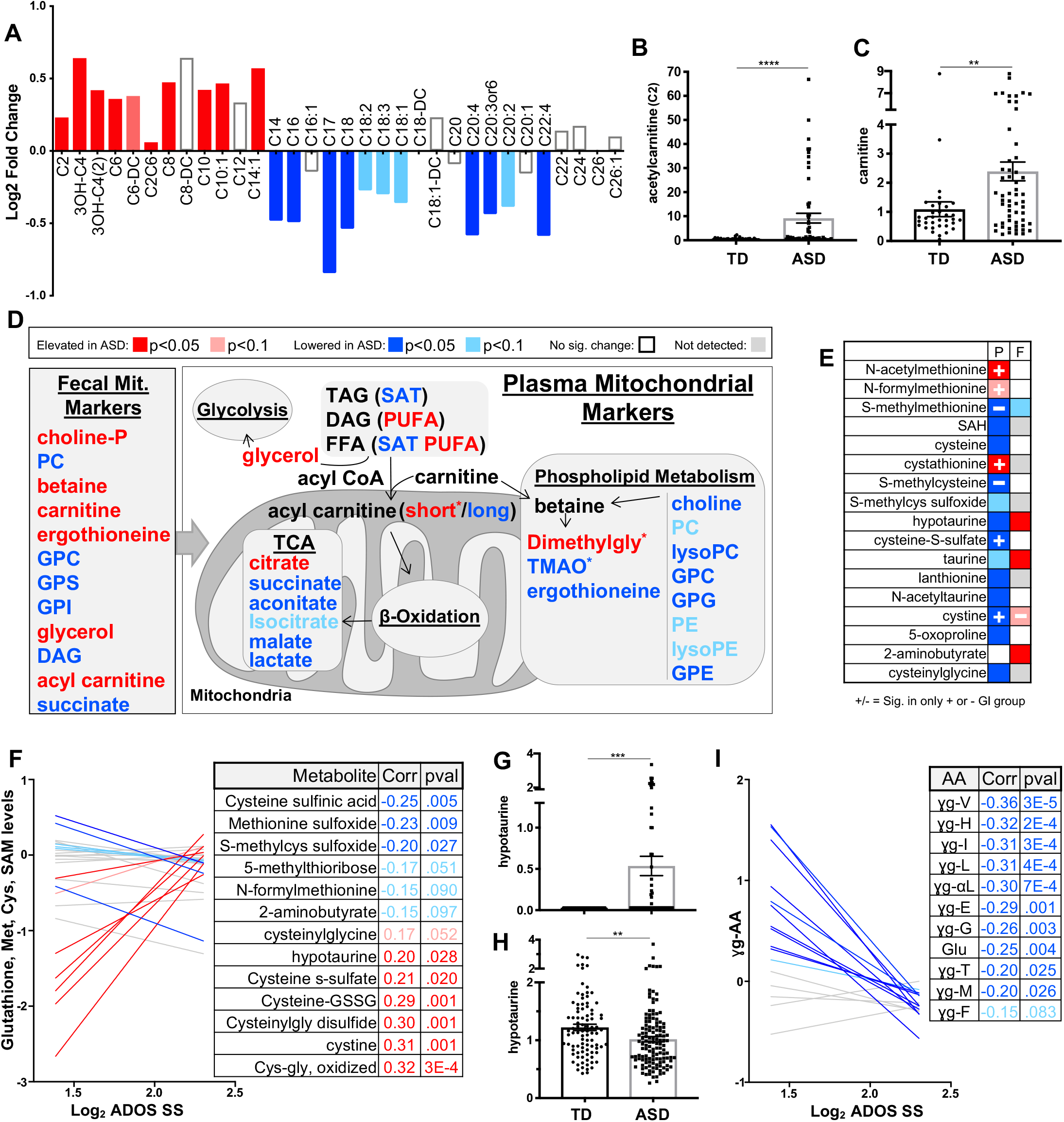
ASD Correlates with Cellular Energy and Oxidative Stress Metabolites. (A) Log2 fold change of acyl-carnitines in the plasma of ASD-GI samples compared to controls. Significance indicated by color according to legend below, determined by ANOVA contrasts. (B-C) Scaled intensity values indicating relative levels of acetylcarnitine(C2) and carnitine, respectively, in ASD fecal samples compared to TD controls (all samples). Data are represented as mean ± SEM. Asterisks indicate significance in ANOVA contrasts performed on total metabolomics dataset. p values: 6.44E-05(B) and 2.55E-03(C). (D) Schematic of mitochondrial markers and other metabolites closely associated with cellular energy in plasma (within center box) and feces (boxed to left). *=significant only in ASD-GI. Color of text indicates direction and significance of change according to legend above. (E) Differences in levels of metabolites from the cysteine, methionine, and glutathione pathways, comparing ASD vs TD. Refer to legends below and in D. (F) Correlation of ADOS-SS with metabolites from the cysteine, methionine, and glutathione pathways. Significant and trending metabolites corresponding to the linear regression in the graph are listed along with Pearson coefficients and p-values. Insignificant correlation values are not listed here. Refer to color legend in D. (G-H) Scaled intensity values indicating relative levels of hypotaurine in feces (G) and plasma (H) (all samples). Data are represented as mean ± SEM. Asterisks indicate significance in ANOVA contrasts performed on total metabolomics dataset. p values: 1.61E-05(G) and 1.35E-3(H). (I) Correlations of gamma-glutamyl amino acids with ADOS-SS, color legend in D, with Pearson coefficients and p-values to the right. PC, phosphatidylcholine; PE, phosphatidylethanolamine; GPG, glycerophosphoglycerol; GPC, glycerophosphocholine; GPE, glycerophosphoethanolamine; GPS, glycerophosphoserine; GPI, glycerophosphoinositol; TAG, triacylglycerol; DAG, diacylglycerol; FFA, free fatty acid; SAT, saturated fatty acid; PUFA, polyunsaturated fatty acid. Data are represented as mean ± SEM. *See also Figure S4 and Tables S1 and S2*.

Amino acid signatures of oxidative stress and mitochondrial function are also present in our dataset. Dysregulated amino acid degradation, homeostasis, and import into the brain have been implicated as a cause of neuronal stress in ASD, and supporting metabolomic data has shown perturbations of various amino acid pathways, such as glutamate, methionine, glutathione, and gamma-glutamyl metabolites (*10, 17, 24, 26, 29, 33, 49, 52, 76*), which exhibit differences as well (Figure 4E). We also demonstrate correlations between pathways of oxidative stress (cysteine, methionine, SAM and glutathione pathways) with the ADOS-SS (Figure 4F). Some of these molecules were found in higher levels in the ASD feces and at lower levels in the ASD plasma, such as hypotaurine (Figures 4E, 4G–4H), which might indicate altered fecal production, excretion or differential uptake into the plasma potentially through varied intestinal permeability. Similar to hypotaurine, levels of its precursor, taurine, were significantly increased in ASD fecal samples, although not altered in plasma (Figure S4E and S4F). Taurine plays many roles throughout the host, and has previously been measured at altered levels in ASD, although with little consensus (*17, 18, 24–26, 33, 65, 71, 77*). Hypotaurine and taurine deficiency has been shown to lead to defects in cell differentiation in the brain (*77*) and their dysregulation could alter neuronal signaling (*78*).

Oxidative stress-related glutathione pathway precursors, gamma-glutamyl amino acids, are relevant to ASD through their influence on levels of neurotransmitters such as gamma-aminobutyric acid (GABA) (*78, 79*), which is widely thought to play a role in ASD. In a recent ASD study, an experimental treatment that led to increased gamma-glutamyl AAs and other redox pathway metabolites correlated with improved behavior metrics in children (*65*). Almost every gamma-glutamyl amino acid is negatively correlated with ADOS-SS (Figure 4I). We also observe perturbations in the urea cycle, which processes the amino group of amino acids for excretion in urine (Figure S4G–S4H). Abnormalities in this pathway can be indicative of altered amino acid degradation observed in ASD and can lead to neurotoxic accumulation of nitrogen-containing compounds in the blood (*80*). Together, these results corroborate and extend a growing body of research into altered mitochondrial metabolism and oxidative stress in ASD.

### Differential Phenolic Xenobiotic Metabolite Levels in ASD

Phenolic metabolites, a diverse structural class comprised of thousands of molecules containing a phenol moiety, come from dietary ingredients or from biotransformation of aromatic amino acids by the gut microbiota. In their free phenolic forms they can be readily absorbed through intestinal tissues; however, microbial modification of these molecules can significantly alter their absorption, bioavailability, and bioactivity (*81*), leading to various benefits or harm to the host (*81, 82*). In fact, altered levels of phenolic molecules have been highlighted in many ASD metabolomic studies (*18, 20–26, 28, 31, 32, 53, 65, 83, 84*). However, a consensus of enrichment or depletion of these metabolites across ASD vs. TD groups has yet to be reached, and most studies have only measured a small subset of this structural class of metabolites.

We observed altered levels of phenolic metabolites belonging to several interrelated pathways, including tyrosine, benzoate, and food component and plant metabolites (Figure 5A). Some molecules, such as homovanillate and tyramine, are involved in neurotransmitter metabolism (*18*). Others include hippurate derivatives, which we measured at lower levels in the ASD plasma samples (Figure 5A). Phytoestrogens such as daidzein, genistein, and equol derivatives are elevated in our ASD plasma samples (Figure 5A). These metabolites have purported health benefits but are also known disruptors of endocrine signaling (*85*). Additionally, we detected altered levels of molecules with structural similarity to para-cresol sulfate, a toxic molecule elevated in the urine of young ASD children (*20, 22*). These include, among others, 4-ethylphenyl sulfate (4EPS), 2-ethylphenylsulfate, cresol derivatives, 4-allylphenyl sulfate, and 4-methylbenzenesulfonate, the latter of which was elevated a remarkable 60-fold in a small subset of ASD samples (Figure 5B–5D, Table S1). Some changes in the levels of these phenolic molecules are also observed in animal models of ASD (*50, 86*). Previously, 4EPS was observed at a 46-fold elevated level in the maternal immune activation mouse model, and daily administration of synthetic 4EPS to wild type mice was sufficient to induce an anxiety-like phenotype (*50*). Here, in ASD plasma samples, 4EPS levels were increased 6.9-fold (Figure 5D). Other phenolic metabolite levels correlate with 4EPS in ASD samples, including 4-acetylphenyl sulfate, a derivative of 4EPS, and others. (Figure 5E, S5A–S5B). These observed alterations in phenolic molecules, combined with mounting evidence from previous studies, suggest that phenolic structural metabolites may play a role in ASD.

**Figure 5.**
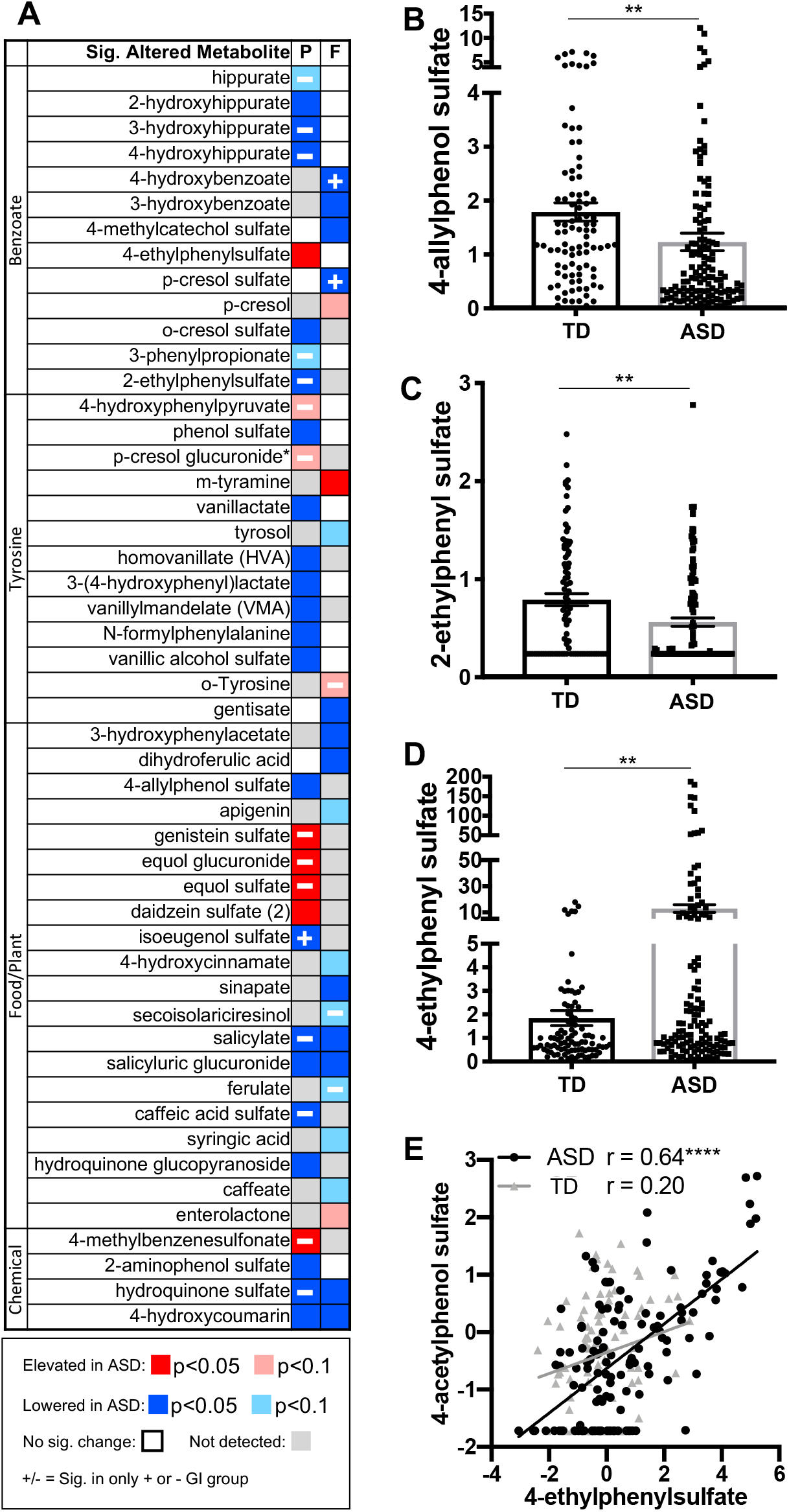
Differential Phenolic Xenobiotic Metabolite Levels in ASD. (A) Phenolic metabolites, belonging to the benzoate, tyrosine, and food component/plant pathways that are significantly different between ASD vs TD groups in plasma (P), and feces (F). Directionality and significance defined in legend. (B) Scaled intensity values indicating relative levels of 4-allylphenol sulfate in plasma (all samples). Asterisks indicate significance in ANOVA contrasts performed on total metabolomics dataset. p value: 2.50-03. (C) Scaled intensity values indicating relative levels of 2-ethylphenyl sulfate in plasma (all samples). (D) Scaled intensity values indicating relative levels of 4-ethylphenyl sulfate (4EPS) in plasma (all samples). Asterisks indicate significance in ANOVA contrasts performed on total metabolomics dataset. p value: 1.16E-03. Data in (B-D) are represented as mean ± SEM. (E) Nearest neighbor correlation between plasma 4EPS and 4-acetylphenol sulfate, log2 scale. Pearson correlation is indicated. p-values: 0.06(TD) and <1E-04(ASD). *See also Figure S5 and Tables S1 and S2*.

## Discussion

Changes in the metabolome have been linked to a number of neurodevelopmental and neurodegenerative disorders (*87, 88*). The current study includes a comprehensive profiling of the metabolites from 231 plasma and 97 matched fecal samples from ASD and TD individuals who have been extensively behavior tested. We observed that specific individual metabolites and metabolic pathways in lipid, amino acid, and xenobiotic metabolism were altered between ASD and TD groups, and that many of these same molecules and pathways correlated to severity of ASD behavioral scores. The core findings are summarized in Figure 6. Various steroid hormone metabolite levels were elevated in ASD samples. PUFA levels, and short chain acyl-carnitines were generally elevated in ASD plasma, while saturated fatty acid levels and long-chain acylcarnitines were decreased, contributing to a general picture of dysregulated cellular energy and oxidative state, with potential connections to mitochondrial dysfunction in ASD. Some of the lipid and xenobiotic metabolites also showed interesting changes within the ASD group when stratified by the presence of GI symptoms. Finally, many phenolic metabolites, derived largely from host and bacterial metabolism of amino acids, plant polyphenols, and other food components were detected at differential levels in plasma and feces between the comparison groups.

**Figure 6.**
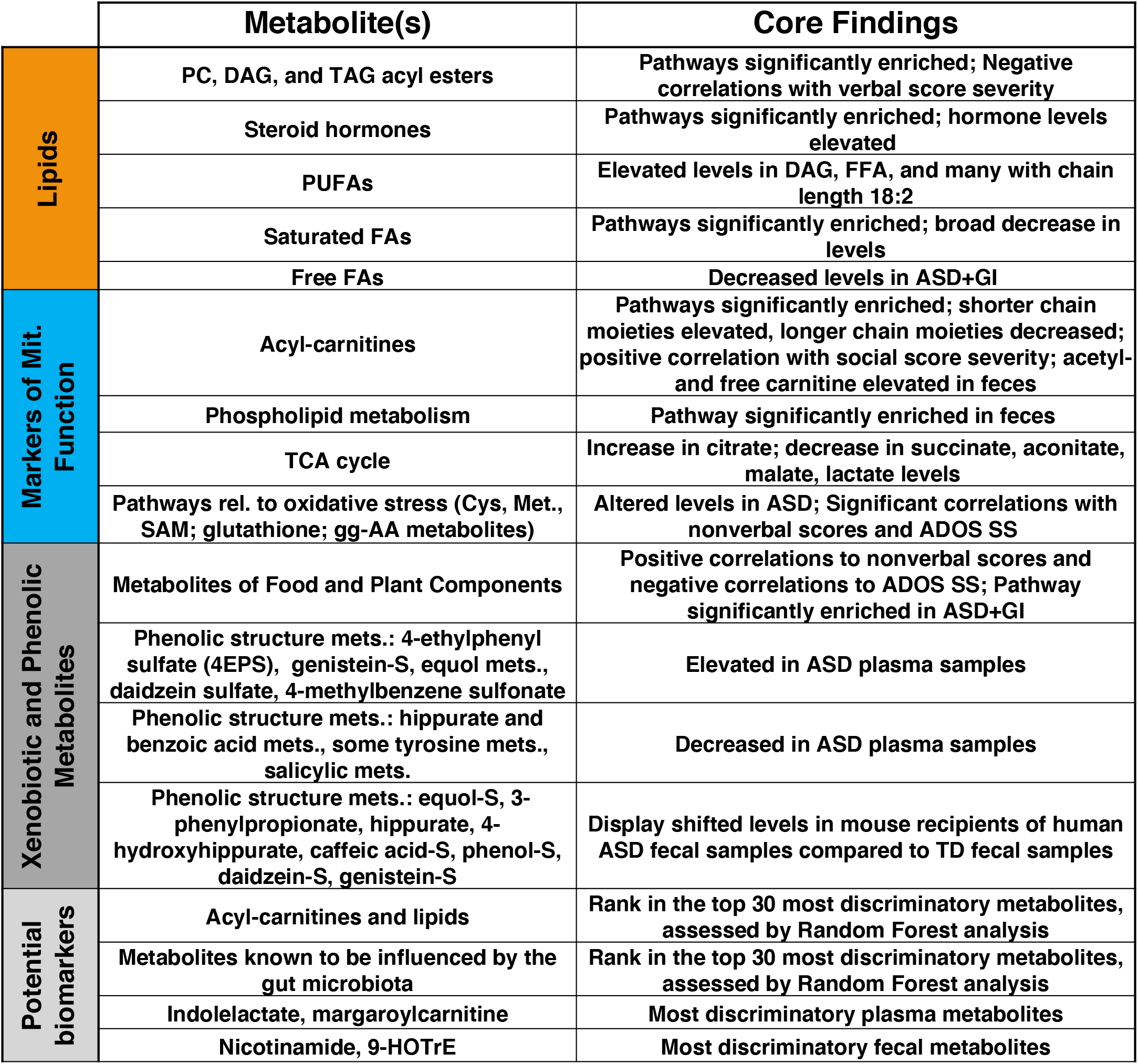
Summary chart of core findings. Categorized into metabolites of lipid, mitochondrial function marker, and xenobiotic and phenolic pathways as well as potential biomarkers, the metabolites of most interest (left) and observations made from the data (right) are summarized.

ASD is diagnosed by behavioral tests, with extensive heterogeneity of symptoms, severity, and etiology between individuals and little consensus on molecular mechanisms. This enigmatic spectrum has a strong but complex genetic basis, with hundreds of reported risk genes (*89*). The contributions of risk alleles to behavior is a vast and active area of research. In addition to genetics, understanding of altered metabolite levels in blood, feces, and brains of ASD individuals may provide a glimpse into physiologic aspects of the disorder and hold the potential to advance diagnosis and/or stratification of sub-populations of ASD.

Interestingly, multiple mouse models support the notion that gut metabolites are associated with brain development and function (*48, 90, 91*). Some metabolites are closely linked with neurological disorders, either to positive or negative outcomes (*92, 93*). Several examples have been reported of gut metabolites entering circulation and directly affecting the brain, as well as cases where metabolites stimulate pathways in the gut, immune, or autonomic nervous system and exert changes to the brain and to behavior (*94–97*). Comprehensive metabolic profiling in humans and animal models provides insight into the molecular status of disease and how genetic factors and environmental risks interact. Deeper analysis of our dataset along with additional studies, with future empirical studies to validate the relevance of our observations, could illuminate aspects of ASD pathophysiology. The intriguing correlations between ASD behaviors, altered levels of fecal and plasma metabolites, and GI symptoms contribute to the concept that ASD may be viewed as a whole-body condition, and argue for increased investigation into peripheral aspects of disease that may lead to advances in diagnosis and improved stratification of ASD populations.

## Acknowledgments

This work was supported by grants from Autism Speaks (Grant #7567 to P.A., A.F. and S.K.M.); the Johnson Foundation (to P.A.); the Brain Foundation (to P.A.); Genome, Environment, Microbiome, and Metabolome in Autism (GEMMA) (#825033 to AF); Axial Biotherapeutics (to S.K.M.); and the National Institutes of Health (HD090214 to P.A.) and (MH100556 to S.K.M.). We would like to thank the MIND Institute study staff, and the dedication and commitment of the families who took part in these studies is gratefully acknowledged.

## SUPPLEMENTAL INFORMATION TITLES AND LEGENDS

**Figure S1.**
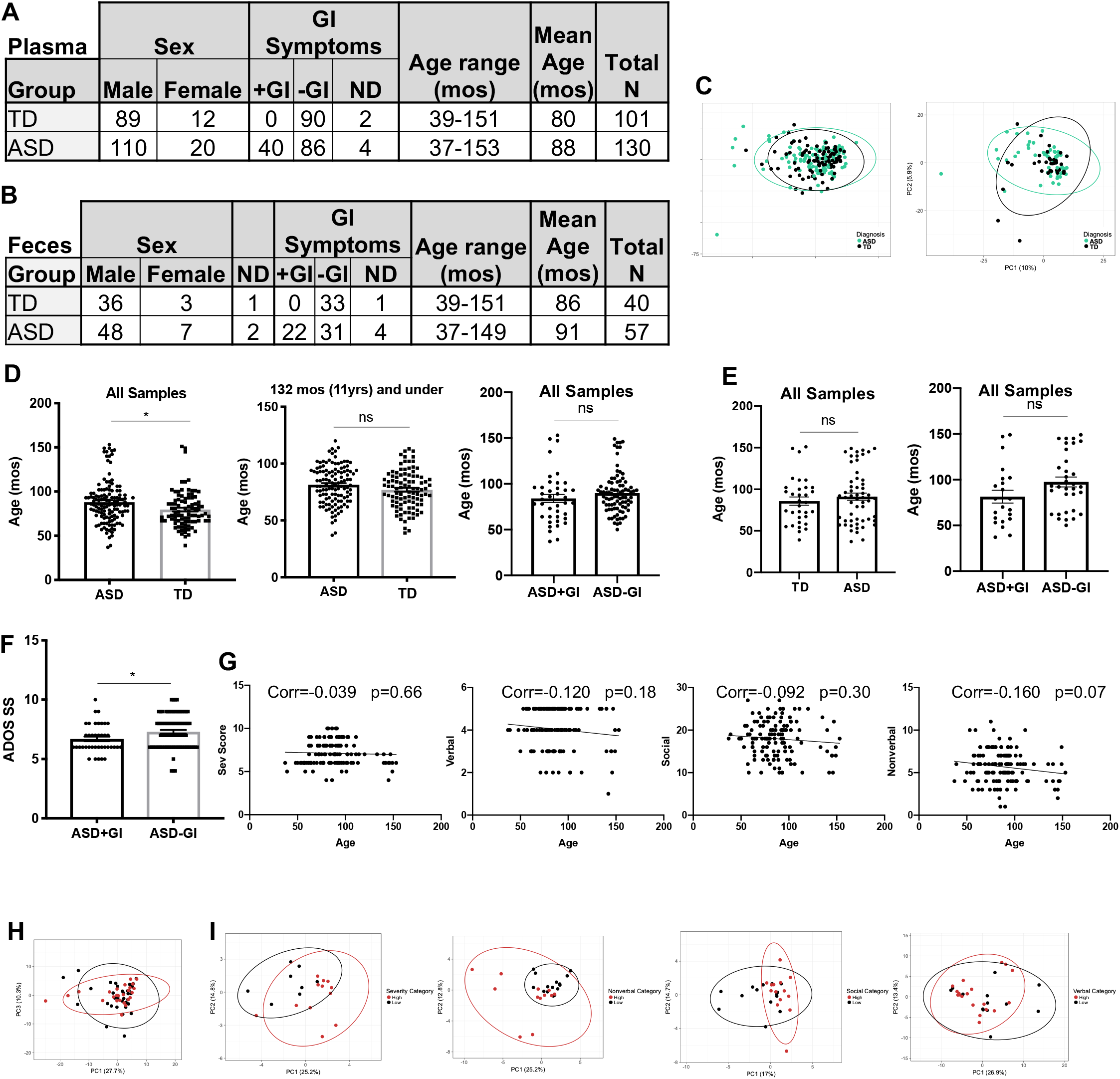
Related to Figures 1 and 2. Group characteristics. (A) Numbers of plasma samples: sex, GI symptoms, age range, and total n. (B) Numbers of fecal samples: sex, GI symptoms, age range, and total n. (C) PCA plot of plasma (left) and feces (right) with all metabolites as input. (D-F) Data are represented as mean ± SEM. (D) Ages of donors of all plasma samples (p=0.007), all plasma samples 11 years and younger (p=0.051), and according to GI status. (p=0.26) (E) Ages of donors of all fecal samples (p=0.15) and samples according to GI status (p=0.075). (F) ADOS-SS graphed according to GI status within the ASD sample group. (p=0.018) (G) Correlations between age (in months) ASD plasma sample individuals and behavior metrics are shown, with linear regression line, spearman correlation value, and p-value. (H) PCA of plasma metabolites significantly correlated with nonverbal behavior, grouped by severity. Ellipses indicate 95% confidence interval. (I) PCA of fecal metabolites significantly correlated with behavior scores, grouped by severity. Ellipses indicate 95% confidence interval.

**Figure S2.**
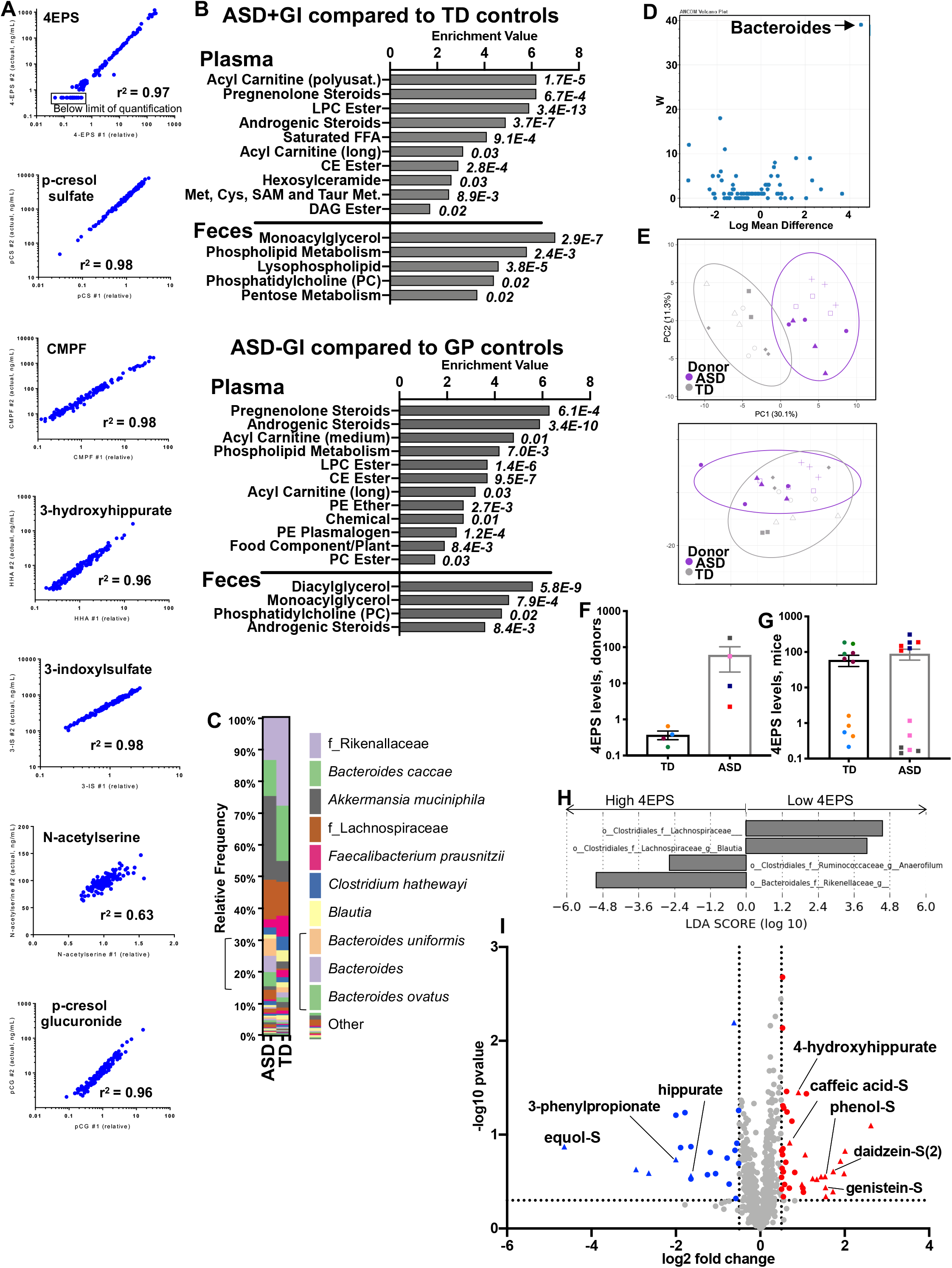
Related to Figures 1 and 2. Relative vs absolute values for select metabolites, pathway enrichment analysis of ASD-GI and ASD+GI comparisons to TD controls, and transfer of human fecal microbiota into mice. (A) Correlation plots comparing the relative abundance measured in the untargeted analysis and the follow-up quantitated values for a subset of 139 plasma samples. Correlations for 4-ethylphenyl sulfate, p-cresol sulfate, 3-carboxy-4-methyl-5-propyl-2-furanpropanoate (CMPF), 3-hydroxyhippurate, 3-indoxylsulfate, N-acetylserine, and p-cresol glucuronide are shown. (B) Significantly affected pathways in ASD+GI vs TD and ASD-GI vs TD comparisons with enrichment value plotted and p-value to the right of each bar. At the top plasma sample results are displayed, with fecal sample results below. (C) 16s profiles for ASD and TD colonized mice. When identified, genus and species are indicated. When family was the most specific identifier, noted by f in legend. (D) Volcano plot for comparison of 16s sequencing in recipient ASD compared to TD mice, genus level. Arrow indicating significant result of *Bacteroides* genus, W value=39. (E) PCA of ASD and TD colonized mice, with all significantly different metabolites from ANOVA contrasts used as input (top) and all metabolites as input (bottom). Ellipses indicate 95% confidence intervals (F) Scaled intensity values indicating relative levels of 4EPS levels in plasma from donors used to colonize mice. p-value 0.23 (G) Scaled intensity values indicating relative levels of 4EPS levels in mice colonized with TD or ASD donors, colored according to donor. Log10 scaled intensity of biochemical peaks is plotted along the y-axis. (H) LeFSe analysis of fecal bacteria at the genus level from mice exhibiting high or low 4EPS levels (regardless of donor diagnosis) after colonization with human samples. (I) Volcano plot of −log10 p-value and log2 fold change of all mouse metabolites analyzed by mixed effects model. All metabolites above a log2 fold change of +/−0.5 and a −log10 p-value of 0.3 are colored. Triangles denote phenolic metabolites. Those phenolic metabolites also significantly altered in the human samples are labeled by name. *See also Table S5*.

**Figure S3.**
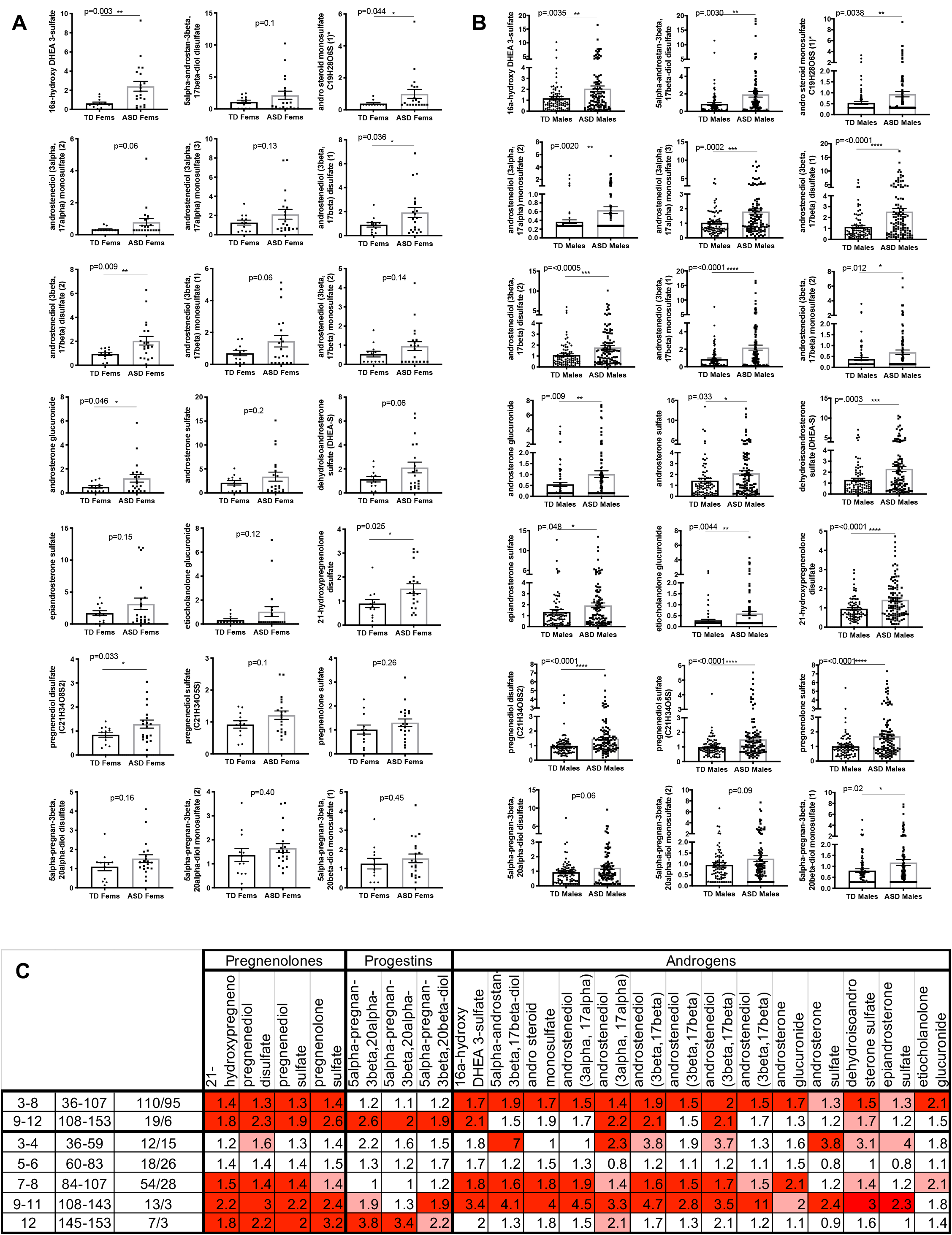
Related to Figure 3. Steroid hormones by sex and age. (A) Levels of androgens, pregnenolones and progestins in all female plasma samples. Scaled intensity values for each female plasma sample. p-values of Welch’s corrected t test are shown about each graph. (B) Levels of androgens, pregnenolones and progestins in all male plasma samples. Scaled intensity values for each male plasma sample shown. p-values of Welch’s corrected t test are shown about each graph. (C) Pregnenolone, progestin, and androgen steroid metabolites clustered by tighter age groupings, denoted in years or months. Dark red indicates significant difference (pval<0.05), light red indicates a trend (0.05>pval>0.1), with fold change in text.

**Figure S4.**
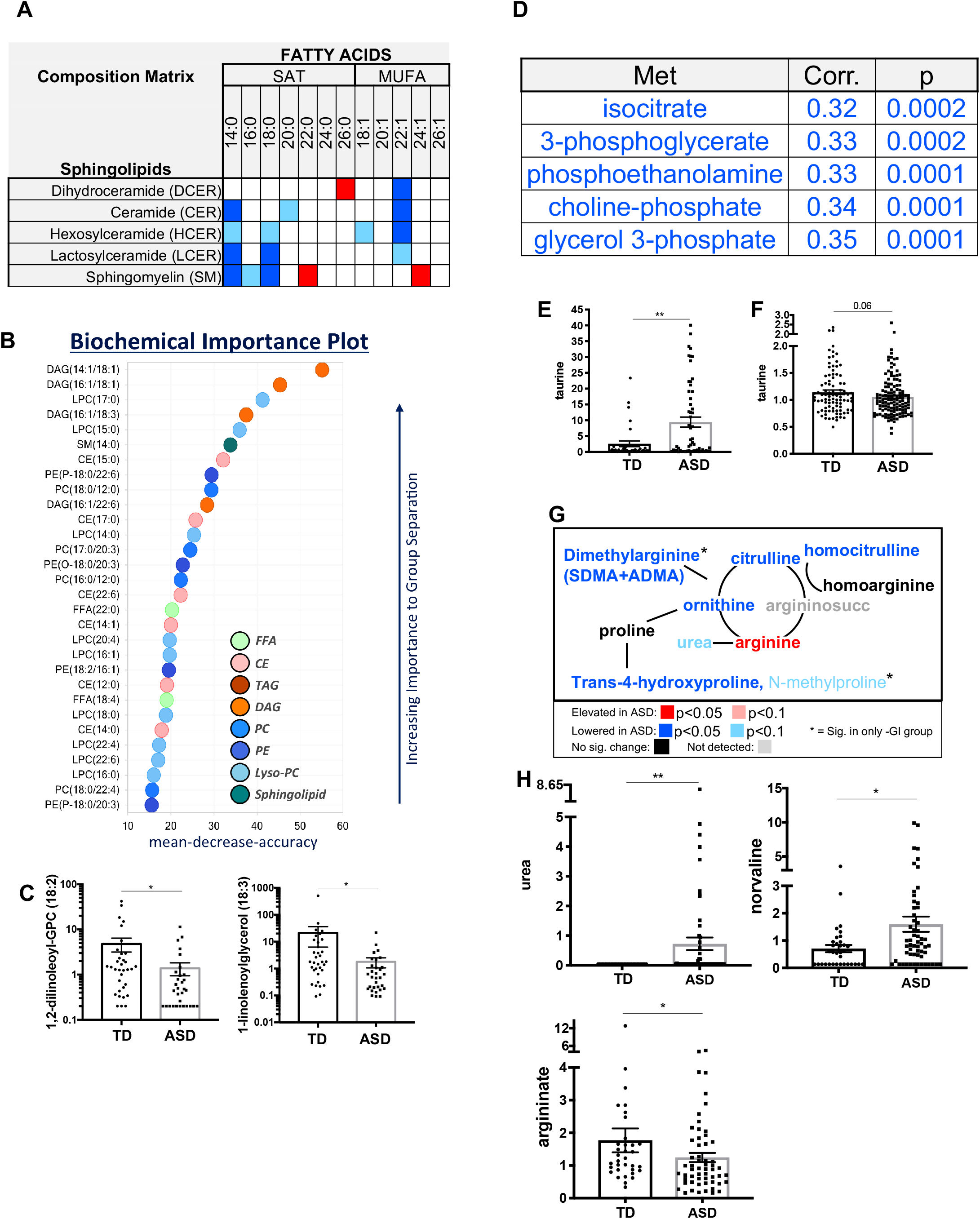
Related to Figures 3 and 4. Complex lipid panel, cellular energy, and amino acid results. (A) Sphingolipids with significantly altered levels in plasma of ASD compared to TD (all samples). (B) Biochemical importance plot of Random Forest analysis using complex lipid panel results for all plasma samples. (C) Scaled intensity values indicating relative levels of two representative PUFA (18:2)-containing fecal lipids that differ in directionality from the plasma phenotype shown in main text Figure 3B. Asterisks indicate significance in ANOVA contrasts performed on total metabolomics dataset. p values: 4.5E-04(left) and 5.9E-04(right). Data are represented as mean ± SEM. (D) The top 5 most positively correlated plasma metabolites with ADOS-SS. The metabolite, spearman correlation value, and p-value are listed. (E) Scaled intensity values indicating relative levels of taurine in feces (all samples). Asterisks indicate significance in ANOVA contrasts performed on total metabolomics dataset. p value: 0.012. Data are represented as mean ± SEM. (F) Scaled intensity values indicating relative levels of taurine in plasma (all samples). p-value from ANOVA contrasts performed on total metabolomics dataset. Scaled intensity of taurine levels in plasma (all samples). (G) Abbreviated pathway schematic of significantly different metabolites in the urea cycle in plasma. Asterisk indicates significance only in the ASD-GI/TD comparison. (H) Scaled intensity values indicating relative levels of urea cycle metabolites that are altered in feces: urea, norvaline, and argininate (all samples). Asterisks indicate significance in ANOVA contrasts performed on total metabolomics dataset. p values: 1.7E-04(urea), 0.02(norvaline), and 0.014(argininate). Data are represented as mean ± SEM.

**Figure S5.**
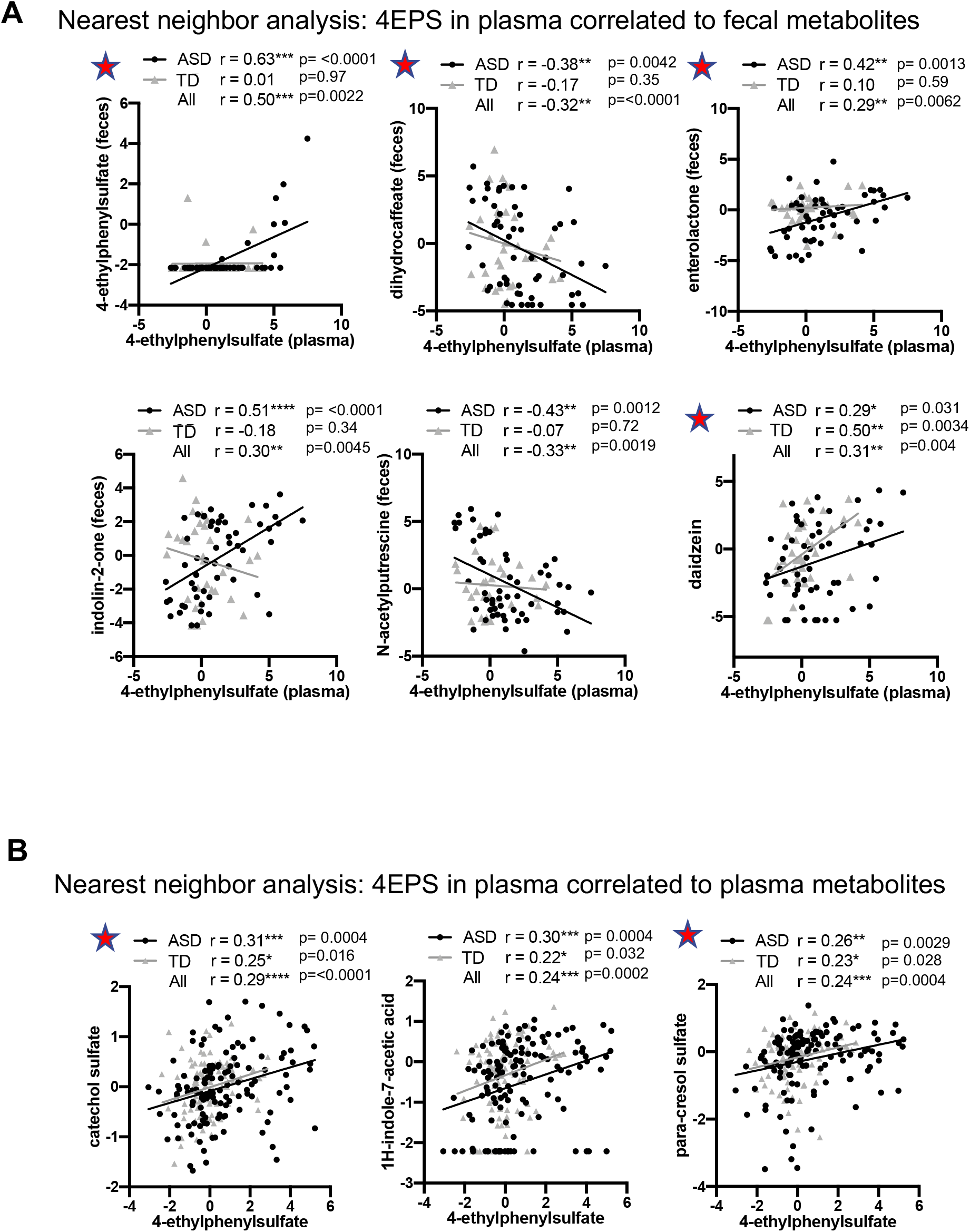
Related to Figure 5. Nearest neighbor analysis of 4EPS. (A) Nearest neighbor analysis of 4EPS plasma levels resulting from correlations with all fecal metabolite levels. Red star denotes phenolic compounds. (B) Nearest neighbor analysis of 4EPS plasma levels against all plasma metabolite levels (minus 4-acetylphenol sulfate from Main Figure 5C). Red star denotes phenolic compounds.

**Table S1. Related to Figures 1–6. Fold change and statistics of human plasma metabolites and correlations to behavior scores. ANOVA contrasts, p-value, q-value, % filled values, and pearson correlation r is provided.**

**Table S2. Related to Figure 1–6. Fold change and statistics of human fecal metabolites and correlations to behavior scores. ANOVA contrasts, p-value, q-value, % filled values, and pearson correlation r is provided.**

**Table S3. Related to Figure 6. Fold change and p value (Welch’s t test) of human donor plasma metabolites.**

**Table S4. Related to Figure 6. Fold change and p value (Welch’s t test) of human donor fecal metabolites.**

**Table S5. Related to Figure 6. Fold change and statistics of mouse plasma metabolites.**

## Supplemental Methods

### Extended Participant Information

Briefly, the GISS consists of 7 sections, each section asking a series of 1 to 6 questions to determine if the participant met the criteria for diarrhea, constipation or irritable bowel syndrome (IBS)-like symptoms, and if stooling/symptoms have been consistent for the last 6 months. Any individual with antibiotic use within 3 months was excluded, but multivitamins and over the counter supplements were permitted. Based on their responses, participants were placed in one of four groups, ASD+GI, ASD-GI, TD+GI or TD-GI. Due to the low incidence of GI issues in typically developing children enrolled in the study, we only have 9 individuals in the TD+GI groups. Our original statistical analysis resulted in many seemingly intriguing metabolites with differential levels between the ASD+GI and TD+GI samples. However, upon closer inspection, most of these differences were driven by the fact that the TD+GI group (with a small n) was very different from all other samples rather than the ASD+GI being unique. There were occasional differences that were indeed convincing between the ASD+GI and TD+GI, but these differences also arose in the comparison of −GI samples, so they were not specific to the GI phenotype and presenting them in an ASD+GI compared to TD+GI context would have been misleading. Thus, the TD+GI group was removed from the analysis and is not presented here.

This study was approved by institutional review boards for the State of California and the University of California, Davis. Informed consent is obtained from a legal guardian for all study participants prior to data collection in accordance with the UC Davis IRB protocol.

### Blood and stool collection

Peripheral blood was collected from participants in acid-citrate dextrose Vacutainers (BD Biosciences; San Jose, Ca). Blood was centrifuged at 2100 rpm for 10 minutes followed by the collection of plasma into cryovials. Plasma was stored at −80 °C until analysis. In addition to blood, stool samples were also obtained, parents were given collection containers with RNA later to collect stool samples at home and asked to store the samples in the freezer and brought back frozen to the clinic within 24 hrs.

### Metabolite analysis

All metabolite analysis, identification, quality control were performed by standard procedures at Metabolon Inc. as follows.

### Sample Preparation

All samples were maintained at −80°C until processed. Samples were prepared using the automated MicroLab STAR^®^ system from Hamilton Company. Several recovery standards were added prior to the first step in the extraction process for QC purposes. To remove protein, dissociate small molecules bound to protein or trapped in the precipitated protein matrix, and to recover chemically diverse metabolites, proteins were precipitated with methanol under vigorous shaking for 2 min (Glen Mills GenoGrinder 2000) followed by centrifugation. The resulting extract was divided into five fractions: two for analysis by two separate reverse phase (RP)/UPLC-MS/MS methods with positive ion mode electrospray ionization (ESI), one for analysis by RP/UPLC-MS/MS with negative ion mode ESI, one for analysis by HILIC/UPLC-MS/MS with negative ion mode ESI, and one sample was reserved for backup. Samples were placed briefly on a TurboVap^®^ (Zymark) to remove the organic solvent. The sample extracts were stored overnight under nitrogen before preparation for analysis.

### QA/QC

Several types of controls were analyzed in concert with the experimental samples: a pooled matrix sample generated by taking a small volume of each experimental sample (or alternatively, use of a pool of well-characterized human plasma) served as a technical replicate throughout the data set; extracted water samples served as process blanks; and a cocktail of QC standards that were carefully chosen not to interfere with the measurement of endogenous compounds were spiked into every analyzed sample, allowed instrument performance monitoring and aided chromatographic alignment. Instrument variability was determined by calculating the median relative standard deviation (RSD) for the standards that were added to each sample prior to injection into the mass spectrometers. Overall process variability was determined by calculating the median RSD for all endogenous metabolites (i.e., non-instrument standards) present in 100% of the pooled matrix samples. Experimental samples were randomized across the platform run with QC samples spaced evenly among the injections.

### Ultrahigh Performance Liquid Chromatography-Tandem Mass Spectroscopy (UPLC-MS/MS)

All methods utilized a Waters ACQUITY ultra-performance liquid chromatography (UPLC) and a Thermo Scientific Q-Exactive high resolution/accurate mass spectrometer interfaced with a heated electrospray ionization (HESI-II) source and Orbitrap mass analyzer operated at 35,000 mass resolution. The sample extract was dried then reconstituted in solvents compatible to each of the four methods. Each reconstitution solvent contained a series of standards at fixed concentrations to ensure injection and chromatographic consistency. One aliquot was analyzed using acidic positive ion conditions, chromatographically optimized for more hydrophilic compounds. In this method, the extract was gradient eluted from a C18 column (Waters UPLC BEH C18-2.1×100 mm, 1.7 μm) using water and methanol, containing 0.05% perfluoropentanoic acid (PFPA) and 0.1% formic acid (FA). Another aliquot was also analyzed using acidic positive ion conditions, however it was chromatographically optimized for more hydrophobic compounds. In this method, the extract was gradient eluted from the same afore mentioned C18 column using methanol, acetonitrile, water, 0.05% PFPA and 0.01% FA and was operated at an overall higher organic content. Another aliquot was analyzed using basic negative ion optimized conditions using a separate dedicated C18 column. The basic extracts were gradient eluted from the column using methanol and water, however with 6.5mM Ammonium Bicarbonate at pH 8. The fourth aliquot was analyzed via negative ionization following elution from a HILIC column (Waters UPLC BEH Amide 2.1×150 mm, 1.7 μm) using a gradient consisting of water and acetonitrile with 10mM Ammonium Formate, pH 10.8. The MS analysis alternated between MS and data-dependent MS^n^ scans using dynamic exclusion. The scan range varied slighted between methods but covered 70-1000 m/z. Raw data files are archived and extracted as described below.

### Bioinformatics

The informatics system consisted of four major components, the Laboratory Information Management System (LIMS), the data extraction and peak-identification software, data processing tools for QC and compound identification, and a collection of information interpretation and visualization tools for use by data analysts. The hardware and software foundations for these informatics components were the LAN backbone, and a database server running Oracle 10.2.0.1 Enterprise Edition.

### LIMS

The purpose of the Metabolon LIMS system was to enable fully auditable laboratory automation through a secure, easy to use, and highly specialized system. The scope of the Metabolon LIMS system encompasses sample accessioning, sample preparation and instrumental analysis and reporting and advanced data analysis. All of the subsequent software systems are grounded in the LIMS data structures. It has been modified to leverage and interface with the inhouse information extraction and data visualization systems, as well as third party instrumentation and data analysis software.

### Data Extraction and Compound Identification

Raw data was extracted, peak-identified and QC processed using Metabolon’s hardware and software. These systems are built on a webservice platform utilizing Microsoft’s.NET technologies, which run on high-performance application servers and fiber-channel storage arrays in clusters to provide active failover and load-balancing. Compounds were identified by comparison to library entries of purified standards or recurrent unknown entities. Metabolon maintains a library based on authenticated standards that contains the retention time/index (RI), mass to charge ratio (*m/z*), and chromatographic data (including MS/MS spectral data) on all molecules present in the library. Furthermore, biochemical identifications are based on three criteria: retention index within a narrow RI window of the proposed identification, accurate mass match to the library +/- 10 ppm, and the MS/MS forward and reverse scores between the experimental data and authentic standards. The MS/MS scores are based on a comparison of the ions present in the experimental spectrum to the ions present in the library spectrum. While there may be similarities between these molecules based on one of these factors, the use of all three data points can be utilized to distinguish and differentiate biochemicals. More than 3300 commercially available purified standard compounds have been acquired and registered into LIMS for analysis on all platforms for determination of their analytical characteristics. Additional mass spectral entries have been created for structurally unnamed biochemicals, which have been identified by virtue of their recurrent nature (both chromatographic and mass spectral). These compounds have the potential to be identified by future acquisition of a matching purified standard or by classical structural analysis.

### Complex Lipids Platform

Lipids were extracted from samples in methanol:dichloromethane in the presence of internal standards. The extracts were concentrated under nitrogen and reconstituted in 0.25mL of 10mM ammonium acetate dichloromethane:methanol (50:50). The extracts were transferred to inserts and placed in vials for infusion-MS analysis, performed on a Shimazdu LC with nano PEEK tubing and the Sciex SelexIon-5500 QTRAP. The samples were analyzed via both positive and negative mode electrospray. The 5500 QTRAP scan was performed in MRM mode with the total of more than 1,100 MRMs. Individual lipid species were quantified by taking the peak area ratios of target compounds and their assigned internal standards, then multiplying by the concentration of internal standard added to the sample. Lipid class concentrations were calculated from the sum of all molecular species within a class, and fatty acid compositions were determined by calculating the proportion of each class comprised by individual fatty acids.

### Peak Quantification, Normalization and Statistical analysis

Peaks were quantified using area-under-the-curve. The present dataset comprises a total of 1611 and 814 compounds of known identity for the human plasma and fecal samples, respectively. The mouse dataset comprises a total of 746 known biochemicals. Following log transformation and imputation of missing values, if any, with the minimum observed value for each compound, ANOVA contrasts using ArrayStudio (Qiagen) were used to identify biochemicals that differed significantly between experimental groups. Analysis by two-way ANOVA identified biochemicals exhibiting significant interaction and main effects for experimental parameters of disease and GI symptoms. For mouse samples, in addition to two-way ANOVA contrasts, in order to account for nonindependence (match to donor), we ran a linear mixed effect model where the donor match is a random effect. The analysis was done in R.gui version 3.5.1. Correlation analysis was performed comparing the relationship between observed changes in metabolite levels and measures of severity. An estimate of the false discovery rate (q-value) is calculated to take into account the multiple comparisons that normally occur in metabolomic-based studies. In all graphs of individual metabolites, the significance noted in the graph is based on p-value from the ANOVA and can be found in the corresponding Supplemental Tables along with the q-values.

### Principle Components Analysis (PCA)

All PCA plots were generated using the ClustVis web tool. In cases where ASD samples were subdivided by severity, samples were separated and grouped by behavioral score as close to quartiles as possible without splitting samples with the same score into separate groups. The groupings were as follows: for human feces scores for lower and higher severity groups, respectively: ADOS scores 4-5 and 8-10; nonverbal scores 1-4 and 7-10, social scores 10-14 and 20-27, verbal scores 1-3 and 5. Groupings for human plasma scores for lower and higher severity groups, respectively: ADOS scores 4-5 and 9-10; nonverbal scores 1-4 and 8-11, social scores 10-14 and 22-27, verbal scores 1-3 and 5.

### Random Forest Analysis

Random Forest Analysis was performed as described previously ^1^. A random subset of the data with identifying true class information was selected to build the tree (“bootstrap sample” or “training set”), and then the remaining data, the “out-of-bag” (OOB) variables, were passed down the tree to obtain a class prediction for each sample. This process was repeated thousands of times to produce the forest. The final classification of each sample was determined by computing the class prediction frequency for the OOB variables over the whole forest. When the full forest is grown, the class predictions were compared to the true classes, generating the “OOB error rate” as a measure of prediction accuracy. The mean decrease in accuracy was determined by randomly permuting a variable, running the observed values through the trees, and then reassessing the prediction accuracy. Thus, the Random Forest analysis provides an “importance” rank ordering of biochemicals.

The further Random Forest using the top 30 metabolites was performed by first leaving out one sample at a time, fitting the Random Forest, keeping the top 30 metabolites, then using the top 30 to predict the observation that was left out. This was repeated for each sample and then the actual result was compared to the predicted and error was calculated.

### Pathway Enrichment Analysis

For each individual pair-wise comparison, pathway enrichment determined the number of statistically significantly different compounds relative to all detected compounds in a sub-pathway, compared to the total number of statistically significantly different compounds relative to all detected compounds in the study. A pathway enrichment value greater than one indicates that the pathway contains more significantly changed compounds relative to the study overall, suggesting that the pathway may be a target of interest for further investigation. Enrichment Value = (# of significant metabolites in pathway(k) / total # of detected metabolites in pathway(m)) / (total # of significant metabolites(n) / total # of detected metabolites(N)) (k/m)/(n/N).

### Animal husbandry and sample collection

All mouse housing and experiments were approved by the California Institute of Technology IACUUC. Fecal samples were thawed, 300mls of 1.5% bicarbonate in PBS was added, samples were vortexed, and then allowed to settle. 150uL of fecal sample was then delivered by oral gavage to 5-week-old germ free mice that were then maintained in sterile, microisolator cages for 3 weeks. After three weeks, fecal samples were collected for 16s sequencing. Blood was collected by cardiac puncture following euthanasia by CO_2_, plasma was isolated on ice using EDTA-treated collection tubes (Thermo), and samples were stored at −80C until analysis. One replicate GP sample was thawed prematurely and thus removed from further analysis.

### Fecal analysis for 16s sequencing

Fecal samples collected and immediately put into empty sterile tubes, flash frozen, and maintained at −80C until processing. Total DNA was isolated with Qiagen DNeasy powersoil extraction kit (Qiagen) following manufacturer instructions. Hypervariable V4 region of the 16s gene was amplified by PCR using 5Xprime master mix (Prime). Barcoded 806 reverse primers and unique forward 515 primer (IDT) were used as previously described. The amplification was confirmed by electrophoresis and the amplified products were purified with Quiaquick PCR purification kit (Qiagen).

Samples were sent to MGH NGS Core facility to be sequenced on the Illumina MiSeq instrument using MiSeq v2 500-cycle sequencing kit, resulting in approximately 25 million paired-end 250 bp reads covering amplicon regions. Data were analyzed QIIME2 software package at the Bioinformatic core at MGH. Low quality score sequencing reads (average Q < 25) were truncated to 240bp followed by filtering using *deblur* algorithm with default setting and the high quality reads were aligned to the reference library using *mafft*. The aligned reads were masked to remove highly variable positions, and a phylogenetic tree was generated from the masked alignment using the FastTree method. Alpha and beta diversity metrics and Principal Component Analysis plots based on Jaccard distance were generated using default QIIME2 plugins. Taxonomy assignment was performed using *feature-classifier* method and naïve Bayes classifier trained on the Greengenes 13_8 99% operational taxonomic units (OTUs). Linear discriminant analysis Effect Size (LEfSe) was performed as described previously^2^ using the Galaxy web application.

